# Adsorption of Pulmonary and Exogeneous Surfactants on SARS-CoV-2 Spike Protein

**DOI:** 10.1101/2022.05.04.490631

**Authors:** Kolattukudy P. Santo, Alexander V. Neimark

## Abstract

COVID-19 is transmitted by inhaling SARS-CoV-2 virions, which are enveloped by a lipid bilayer decorated by a “crown” of Spike protein protrusions. In the respiratory tract, virions interact with surfactant films composed of phospholipids and cholesterol that coat lung airways. Here, we explore by using coarse-grained molecular dynamics simulations the physico-chemical mechanisms of surfactant adsorption on Spike proteins. With examples of zwitterionic dipalmitoyl phosphatidyl choline, cholesterol, and anionic sodium dodecyl sulphate, we show that surfactants form micellar aggregates that selectively adhere to the specific regions of S1 domain of the Spike protein that are responsible for binding with ACE2 receptors and virus transmission into the cells. We find high cholesterol adsorption and preferential affinity of anionic surfactants to Arginine and Lysine residues within S1 receptor binding motif. These findings have important implications for informing the search for extraneous therapeutic surfactants for curing and preventing COVID-19 by SARS-CoV-2 and its variants.

---

The COVID-19 pandemic, that affected over 2 years more than half-a-billion people and claimed more than six million deaths around the world, has triggered broad research activities aiming at preventing and curing coronavirus disease. COVID-19 is transmitted by inhaling airborne SARS-CoV-2 virus particles, which infect the respiratory system and cause severe acute respiratory syndrome (SARS) that can lead to lung failure. Whereas our knowledge of the biochemical structure and functions of SARS-CoV-2 is quickly growing, the physico-chemical aspects of virus interactions with the respiratory system environment have been sparsely addressed and are poorly understood. Bridging this knowledge gap is important for informing clinical studies on surfactant therapies by administering respiratory and exogenous surfactants that can hinder virus proliferation and penetrations into the cells. The search for suitable therapeutic surfactants is actively expanding.^1–4^ Here, we explore by using coarse-grained molecular dynamics (CGMD) simulations the mechanisms of preferential adsorption of typical pulmonary and exogenous surfactants on SARS-CoV-2 virions that may lead to virus inactivation.

SARS-CoV-2 virions, which contain and protect the virus RNA, are spheroidal nanoparticles, with a lipid bilayer envelope of about 85 nm diameter decorated by a “crown” of 20 nm long protrusions of Spike protein. Being inhaled, SARS-CoV-2 virions travel through lung airways and interact with respiratory surfactant monolayer films coating alveoli that composed of phospholipids, cholesterol, and surfactant proteins, known as the lung surfactant (LS). The virion attachment to cells occurs by binding of S1 domain of Spike protein to angiotensin converting enzyme 2 (ACE2) receptor on the type II pneumocytes of the alveoli.^5–6^ The LS film regulates surface tension at the alveolar interfaces to help breathing and prevent lung collapse during expansion-contraction cycles.^7^ LS film is also the first line of defense against inhaled airborne particles.^3^ Surfactant-induced inactivation of viruses has been a traditional topic of clinical studies before the COVID-19 pandemic.^4, 8–13^ Surfactants, that have been found effective in inactivating various types viruses and in restoring the lung damage, include soap^10^, sodium dodecyl sulfate (SDS),^11^ phospholipids,^12^ sodium laureth sulphate,^13^ and potassium oleate^9^ (see review by Simon et al^4^). The virus inactivation may occur (1) by surfactant penetration into viral envelope lipid membrane leading to its lysis and (2) surfactant adsorption on viral proteins that plays crucial role in the infection process and the stability of the virus.^4^ While viral lysis was observed at high surfactant concentrations that can be cytotoxic, functional inactivation of viral proteins by surfactant adsorption was found at low surfactant concentrations suitable for therapeutic administration.^4^

Despite recent great progress in molecular simulations of coronaviruses, including simulations of systems containing several to hundreds of millions of atoms,^14–16^ no attempts have been made to study the interfacial interactions of SARS-CoV-2 virions with pulmonary and exogenous surfactants that is the focus of this work. Our main goal is to mimic *in-silico* surfactant adsorption on the S1 domain that is responsible for ACE2 binding. Spike protein, a trimer of the 1273-residue S-protein,^17^ has two functional domains, S1 (1-685) and S2 (686-1273). S1 binds to ACE2 with its receptor binding domain (RBD 319-546), while S2 assists in fusion of the virion and host cell membranes. The N-terminal domain (NTD 14-305) of S1 binds to sugar molecules.^18^ The main functional region in RBD is the receptor binding motif (RBM 438-506).^19^

We perform CGMD MARTINI^20–21^ simulations of surfactant adsorption on the S1 domain. Three typical surfactants are considered: zwitterionic phospholipid dipalmitoyl phosphatidyl choline (DPPC) and cholesterol (CHOL), that are main components of LS films, and anionic sodium dodecyl sulphate, that is considered as a potential exogenous surfactant with a therapeutic effect.^22^ The MARTINI model of S1 domain is constructed utilizing the Spike protein structure predicted by I-TASSER^23–24^ (Figure S1a). Simulations are performed at the physiological conditions (310 K, 1 bar). The S1 domain is initially placed in the center of the 18 nm cubic simulation box filled with water and surfactant molecules dispersed around the protein at random positions. To demonstrate the difference in surfactant – protein interactions, we explore 7 surfactant compositions: 3 single component solutions and 4 binary mixtures containing DPPC and CHOL at 4:1 and 1:1 ratios, DPPC and SDS at 1:1 ratio, and CHOL and SDS at 1:1 ratio. The total number of surfactant molecules *N_surf_* varies between 10-50. 4 replicas of each system are simulated for good statistics (in total - 136 simulations). The simulations are run for 1*μs* and the statistics is averaged over the last 200 ns and over the replicas. Details of the CG models schematically shown in Figure 1 and CGMD simulations are provided in the Supporting Information (SI) Section 1.

**Figure 1.**
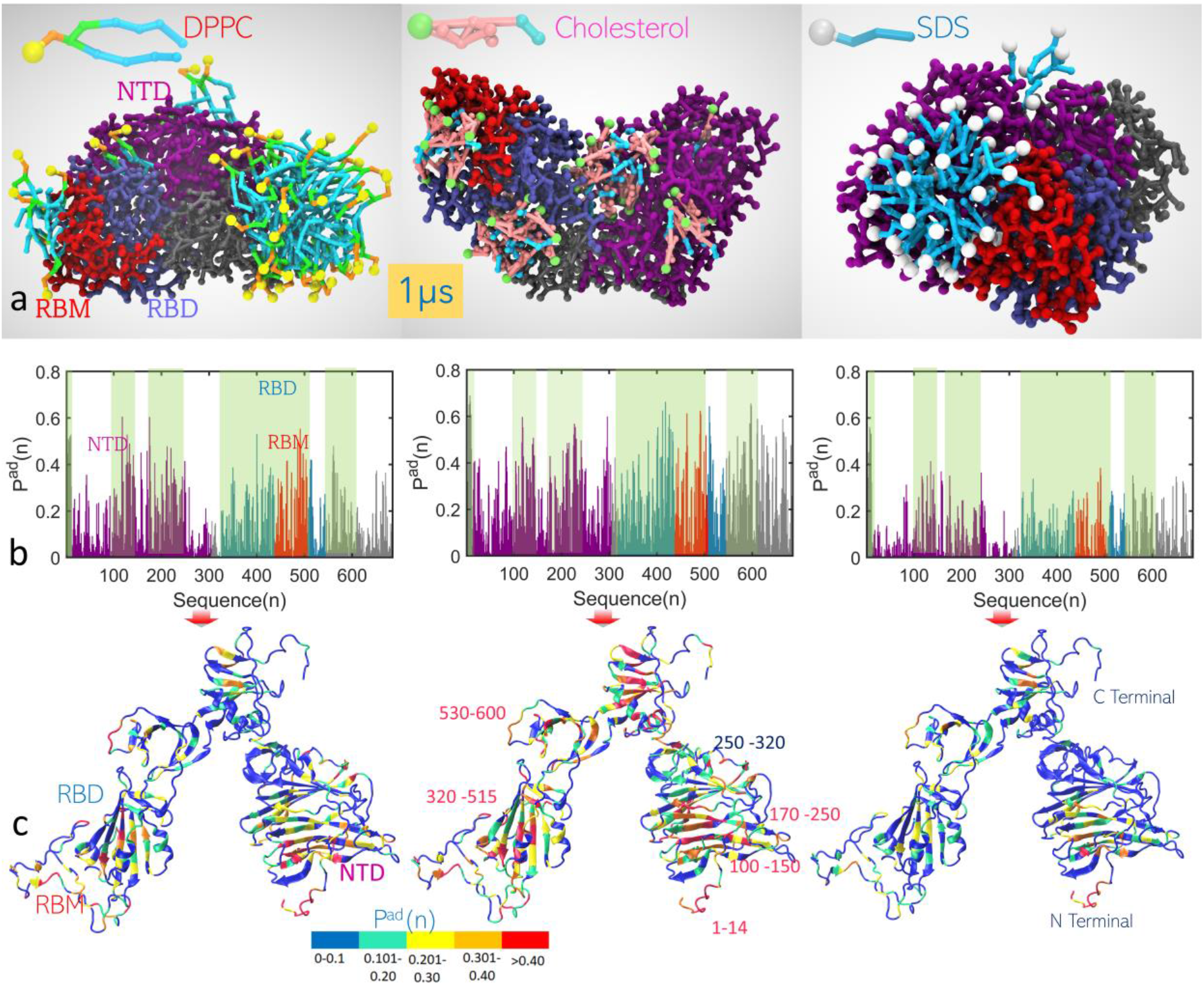
a) Representative snapshots of the surfactant-adsorbed S1 sub-domains of Spike protein: DPPC (left), CHOL (middle) and SDS (right). S1 colors: NTD - purple, RBM - red, RBD - ice blue, other S1 regions - gray. The CG models of surfactants are shown in left upper corners. b) The surfactant adhesion probability of residues along the S1 sequence. The highlighted (light-green) regions correspond to high adsorption probabilities. c) The S1 atomistic structure with residues colored according to the surfactant adsorption probability.

We find that the differences in sorption of various surfactants on S1 domain are determined by an interplay of surfactant-surfactant and surfactant-protein interactions. The former interactions cause surfactant aggregation (micellization). In the single component surfactant systems (Figure 1a), DPPC shows a high tendency to aggregate into large clusters that adhere to the protein, which results in fewer direct lipid-protein contacts. In comparison, cholesterol exhibits a much lower aggregation tendency owing to its wedge-like shape that hinders micellization and, consequently, enhances direct lipid-protein interactions. SDS forms pronounced micellar aggregates, similarly to DPPC. Being a smaller molecule with a simpler structure, SDS adsorbs to the protein more effectively compared to DPPC (see below). All three surfactants show strong affinity to specific S1 functional subdomains, NTD, RBD and RBM.

Preferential adsorption of surfactants on the amino acid residues along the S1 residue sequence is quantified by the probability, *P^ad^*(*n*), of the *n*-th amino acid residue along the sequence (*n* = 1,.., 685) to be in direct contact with adsorbed surfactant. *P^ad^*(*n*) is calculated as the fraction of time that given S1 residue is within 0.62 nm distance from a surfactant molecule (SI, section 2). Note that here we deal with reversible physical adsorption of surfactants. *P^ad^*(*n*) is averaged over the data from 4 replicate simulations with 5 different surfactant concentrations. Averaging over a total of 20 simulations for each surfactant provides reliable statistics for determining the regions of the protein that are preferential for surfactant adsorption. As depicted in Figure 1b-c, *p^ad^*(*n*) exhibits similar patterns for all the three surfactants, which identify the blocks of residues,1-14, 100-150, 170-250, 320-506 and 530-600, as having high affinity to surfactants, in contrast to blocks 14-100, and 250-320, where surfactant adsorption is less probable (see detailed analysis in Figure S3-S4). Figure 1c demonstrates the distribution of S1 residues with respect of their affinity to surfactant adsorption. Note that the RBD and RBM sub-domains represent preferential regions for adhesion of surfactant aggregates. The difference in adsorption of the considered surfactants is further demonstrated with the overall surfactant adsorption probability 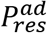 averaged across the whole S1 domain (SI section 2) at a given molar surfactant concentration, that predicts distinctively a higher adsorption probability of cholesterol compared to DPPC and SDS, 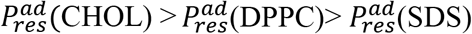 (Figure 2a).

**Figure 2.**
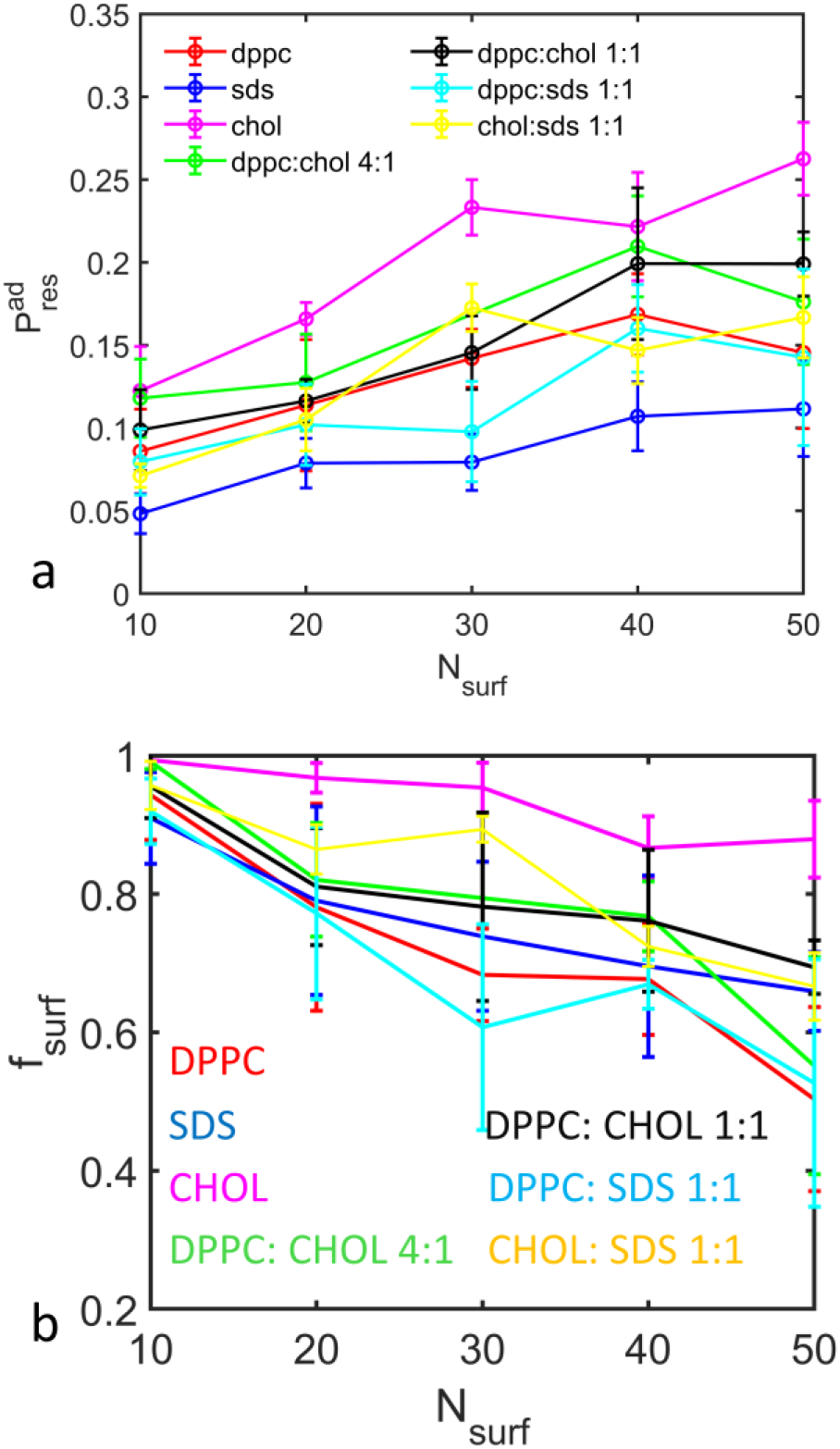
a) The overall probability of surfactant adsorption to S1 domain 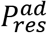 in the single component surfactant solutions and their binary mixtures as a function of surfactant concentration. b) The fraction of surfactant molecules being in direct contact with the protein for different surfactant compositions. Error-bars indicate deviation over 4 replicas.

The fraction of surfactant molecules that are directly adsorbed on the protein, *f_surf_*, indicates the extent of protein-surfactant interactions. Stronger surfactant-surfactant interaction induces formation of larger aggregates and a smaller number of direct lipid-protein contacts. *f_surf_* decreases as the surfactant concentration increases due to surfactant aggregation (Figure 2b), as not all the surfactant molecules in the aggregate are in direct contact with the protein. *f_surf_* follows the order *f_surf_* (CHOL)> *f_surf_* (SDS) > *f_surf_* (DPPC), indicating distinctively higher cholesterol adsorption. Note that SDS has a higher affinity to S1 compared to DPPC, despite lower adsorption probabilities (Figure 1b-c and 2a). SDS is a much smaller molecule and thus make less direct contacts with residues compared to DPPC.

The fractions of different surfactant fragments (heads and tails) or a bead type that are in contact with the protein, *f_bead_* provides a more detailed characterization of the adsorbed surfactants (Figure 3). For instance, *f*_*PO*_4__ represents the ratio of number of PO4 beads in direct contact with the protein residues to the total number of PO4 beads, and so on. In one-component surfactant systems (Figure 3b), *f_bead_* reveals: *f_bead_*(CHOL) > *f_bead_*(SDS) > *f_bead_* (DPPC), which is a much-pronounced trend compared to *f_surf_*. Adsorption of DPPC occurs preferentially by hydrophobic tail and glycerol backbone. There is a preference for adsorption of the anionic PO4 over the cationic choline that has the least affinity to the protein. In SDS, the anionic SO_3_ group has a like affinity to the protein as other uncharged fragments. In general, a clear preference for adsorption of anionic fragments is observed in all systems considered. For cholesterol, the highest affinity to the protein is found for the hydrophilic ROH fragment followed by the aromatic R1-R5 and the aliphatic tail C1/C2 fragments. Note that the observed difference in surfactant adsorption correlates with the affinity of the surfactant head groups. This is explained in part by the fact that in surfactant aggregates adhering to the protein, direct contacts of the tail groups are hindered, while the head groups are located closer to the protein.

**Figure 3.**
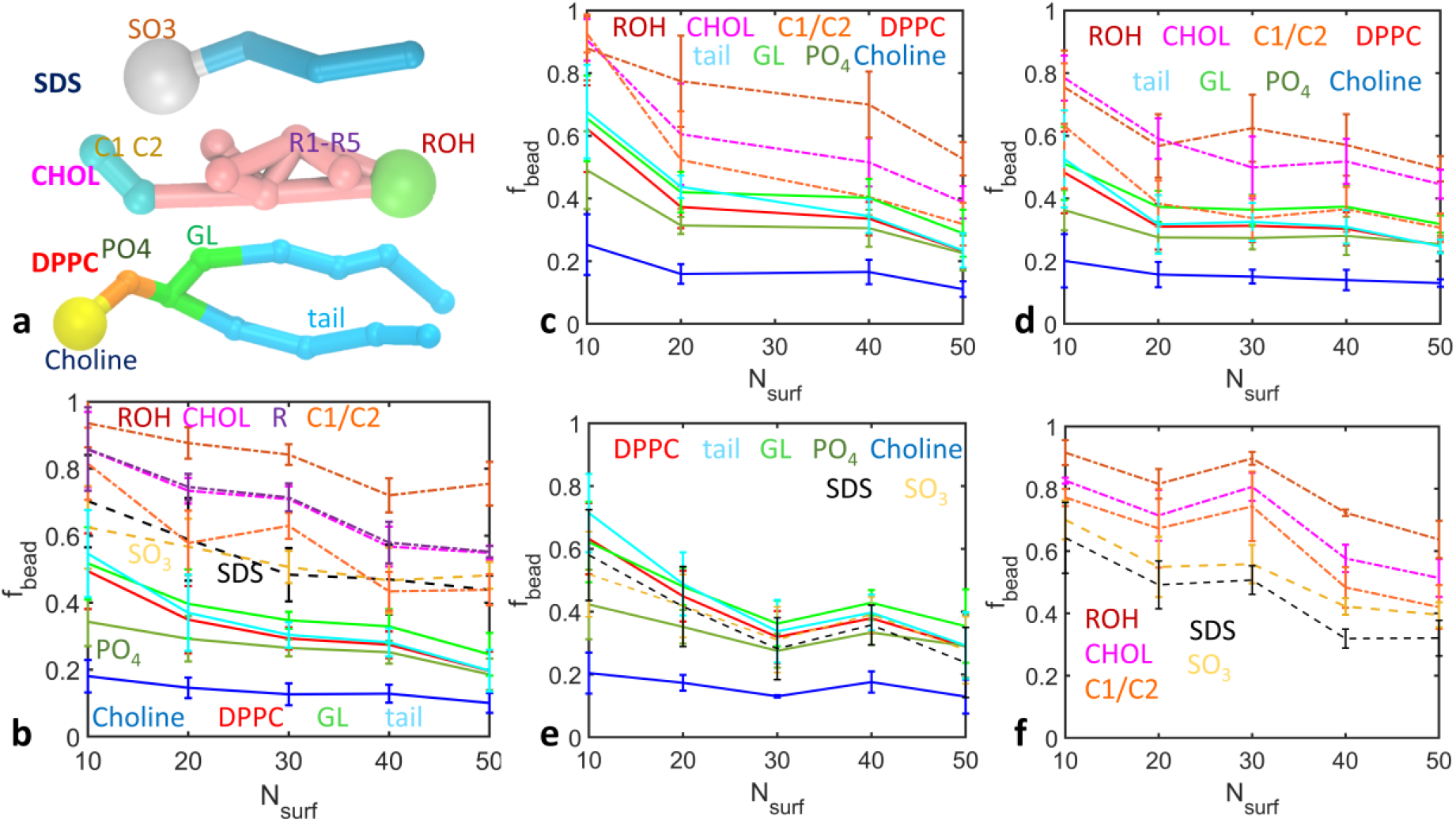
a) Coarse-grained models of surfactant molecules. DPPC consists of the cationic choline, anionic phosphate (PO_4_), glycerol and hydrophobic tail beads. Cholesterol has the hydrophilic OH containing ROH bead, beads R1-R5 representing the aromatic ring and the aliphatic tail beads C1-C2. SDS consists of anionic SO_3_^−^ bead and the 3-bead tail. (b-f) Fraction of various surfactant CG bead types in contact with S1 domain in (b) single component surfactant systems and binary mixtures (c) DPPC:CHOL 4:1 (d) DPPC:CHOL 1:1, (e) DPPC:SDS 1:1 and (f) CHOL:SDS 1:1, at different surfactant concentrations. In each plot, the legends indicate with lines of respective colors.

In mixtures, surfactant adsorption is affected by the interactions between different components. In DPPC-CHOL mixtures (Figure 3c-d), cholesterol adsorption is reduced, while DPPC adhesion is slightly enhanced compared to the respective single-component solutions. This is explained by CHOL association with DPPC that hinders the CHOL-protein contacts and reduces the aggregate size, enhancing DPPC-protein adsorption. In DPPC-SDS systems, SDS adsorption is suppressed (Figure 3e) due to the electrostatic attraction between 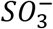 and choline groups that favors DPPC-SDS aggregation. Adsorption in SDS-CHOL systems seem to be unaffected by the inter-surfactant interactions (Figure 3f), although the 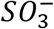 adsorption appears to be somewhat enhanced.

The surfactant affinity to different protein residues is analyzed by calculating the fraction of residues of given type occupied by adsorbed surfactant molecules, *f^ad^*(*J*), where *J* = A,G, …, E, the amino acid residues (SI, section SIII); i.e., *f^ad^*(*A*) = 0.2 means that 20% of the total Alanine residues are occupied. The distribution of different amino acids in S1 domain is given in Figure 4a, and the respective distributions of *f^ad^*(*J*) in Figure 4b. Interestingly, the distributions *f^ad^*(*J*) exhibit the same patterns as shown in Figure 4b independent of the surfactant concentration (Figure S5). Among the hydrophobic residues, which are present in large quantities, ILE, LEU, PHE, PRO and VAL have a higher affinity to surfactants compared to ALA and GLY. Among the polar residues, Cystine and Tyrosine are distinctively preferred by surfactants while abundant ASN, SER and THR are not liked by surfactants.

**Figure 4.**
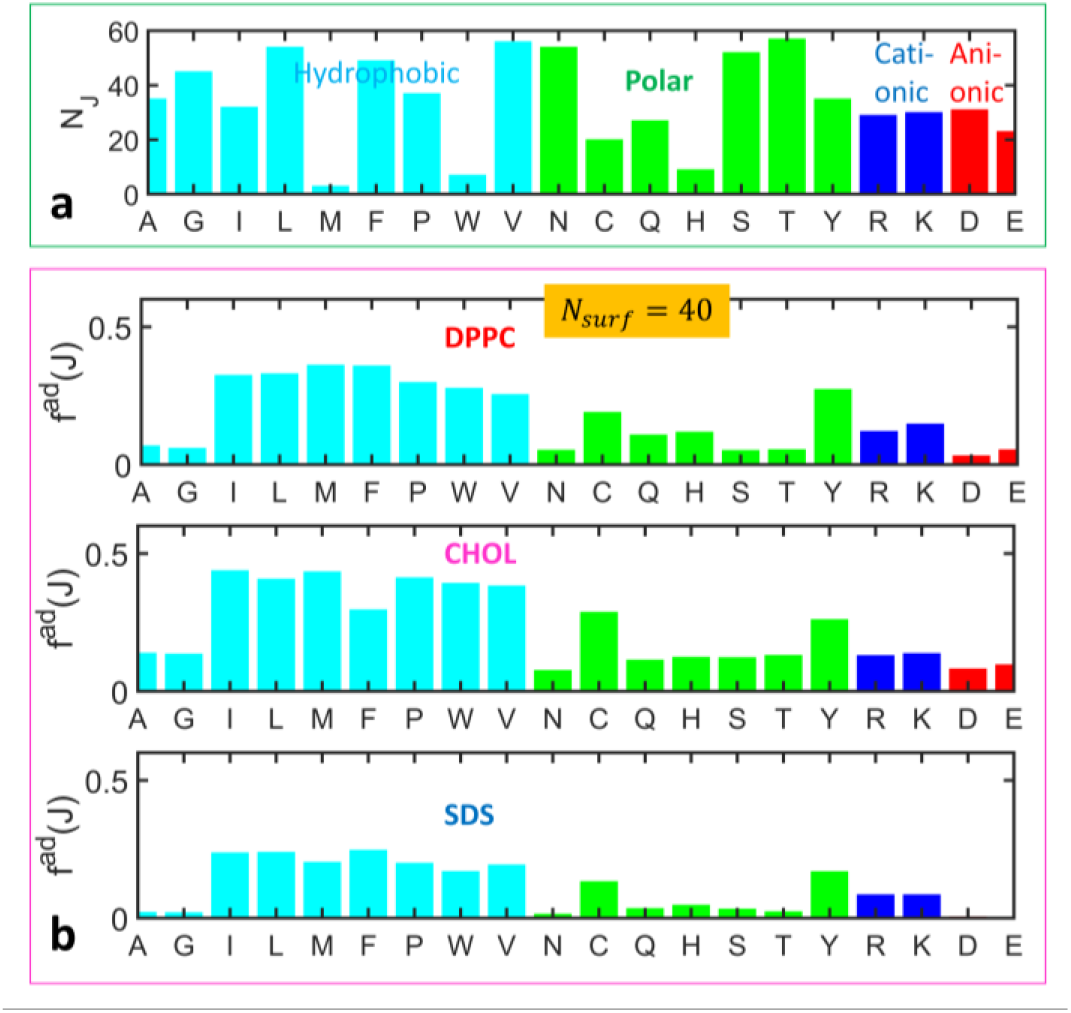
a) The distribution of different amino acid residues in S1. b) Fraction of surfactant adsorbed residue types (*f^ad^*(*J*)) in DPPC, cholesterol and SDS systems at *N_surf_* = 40.

Cationic residues, Arginine and Lysine, are preferred over anionic residues ASP and GLU by all surfactants considered, not just by zwitterionic DPPC and anionic SDS, but also by nonionic cholesterol. These ionic residues, a total of 113, present in similar amounts evenly distributed along the chain, gives the S1 domain an overall cationic character. The observed ability of surfactants to be adsorbed on cationic residues presents a special interest, as the overall cationic character of S1 domain with a net charge of +5 is the driving force for the Spike protein binding to highly negatively charged ACE2 protein.^25^

Understanding the surfactant affinities of various amino acid residues shown in Figure 4 helps to predict the behavior of the CoV-2 variants. S1 domain is strongly cationic with ARG and LYS being preferential sites for surfactant adsorption. However, the roles of ARG and LYS might be not just electrostatic but also residue-specific with additional surfactant interactions with the side chains, which are attractive to non-ionic and zwitterionic surfactants. Noteworthy, the key mutations in the S1 domain of the highly infectious Delta^26^ and, especially, Omicron^27^ variants (Table S2) increase the total number of Arginine and Lysine. The S1 mutations in Delta increase the numbers of ARG by 2 and LYS by 1 and reduce number of GLU by 1, while in the S1 domain of Omicron, ARG is increased by 2 and LYS by 3. The increase in cationic residues may be one of the factors of the high infectivity of CoV-2 variants. In this respect, it is instructive that our simulations suggest that the cationic residues can be effectively targeted by surfactants.

To conclude, our simulations demonstrate that typical zwitterionic and anionic surfactants and cholesterol are strongly adsorbed to the functional fragments of the SARS-CoV-2 Spike S1 domain. The adsorption process proceeds by forming surfactant aggregates adhered on the protein. Adsorption of cholesterol is more efficient than other surfactants, that explains recent experimental observations that cholesterol enhances the Spike interactions with model membranes.^28^ The anionic surfactants have a higher affinity to S1 domain, tending to be adsorbed on cationic Arginine and Lysine residues, which are present in higher quantities in Delta and Omicron variants. The latter effect may have important implications for the search of extraneous therapeutic surfactant that can potentially hinder binding of Spike proteins with ACE2.

## ACKNOWLEDGMENT

Research reported in this publication was supported by the New Jersey Alliance for Clinical and Translational Science (NJACTS) and National Center for Advancing Translational Sciences (NCATS), a component of the National Institute of Health (NIH) under award number UL1TR003017. The content is solely the responsibility of the authors and does not represent the official views of the National Institutes of Health.

The authors thank Andrew Gow and Jared Radbel for useful discussions.

## Supporting Information

### SI. Methods

#### SI. I. Materials and systems

The SARS-CoV-2 Spike protein trimer model (Figure S1a) in the closed state is obtained from the I-TASSER website (https://zhanggroup.org//COVID-19/, lineage A) from which the S1 domain (residues 1-685) is cut. This S1 fragment is converted to MARTINI CG representation (Figure S1a) and the corresponding force field parameters are obtained, using the *martinize.py* script. The CG MARTINI^1–3^ employs a general 4-1 mapping; approximately, 4 heavy atoms are grouped into a CG bead. In the all-atom representation, the S1 domain consist of 10681atoms (including hydrogens) while in the CG model it consists of just 1556 CG atoms. The CG models of the surfactants (Figure S1b), DPPC, Cholesterol and SDS, are taken from the structure files available from the MARTINI website (http://cgmartini.nl/). DPPC is modelled into a 12-bead representation with head group containing the choline, and phosphate beads and two glycerol beads, and two tails containing 4 beads each. Cholesterol has a head group with OH containing ROH group connected to the 5-bead aromatic ring region (R1-R5) and a 2-bead aliphatic tail. SDS is dissociated in the solutions into DS^-^ and sodium ions. The DS^-^ is modelled as consisting of the anionic SO3^-^ and a 3-bead aliphatic hydrophobic tail. The standard MARTINI water model is used as solvent, in which the CG water bead represents 4 water molecules.^3^

GROMACS 2019^4^ versions are used for performing simulations. The systems are constructed with the S1 domain at the center of the simulation box and the surfactants placed distant from each other using the *gmx solvate* command. We simulated systems containing single surfactant components and their binary mixtures; DPPC-, Cholesterol-, and SDS-only systems and DPPC: Cholesterol 4:1, DPPC: Cholesterol 1:1, DPPC: SDS 1:1 and Cholesterol: SDS 1:1 mixtures. The number of surfactants *N_surf_* is varied as 10, 20, 30, 40, and 50, and a total of 34 systems are constructed. The systems are solvated with MARTINI standard water. The S1 domain has a net charge of +5 and to neutralize this charge, 5 Cl-ions are added. In SDS systems, since the SO3 Na head group is ionized and dissociated, and therefore, Na+ ions equal to *N_surf_* are added as well (Table S1). Then four replicas of each system are constructed with a different surfactant placement and the subsequent solvation with water, to improve statistics of the analyzed data, making it a total of 136 systems. Note that since each replica at the same *N_surf_* are constructed independently, the number of water molecules *N_w_* in each replica may differ by a few (~5). The initial system size is 18 × 18 × 18 *nm*^3^ and the typical number of water CG beads is ~49400. Details of the systems are provided in Table S1.

**Figure S3.**
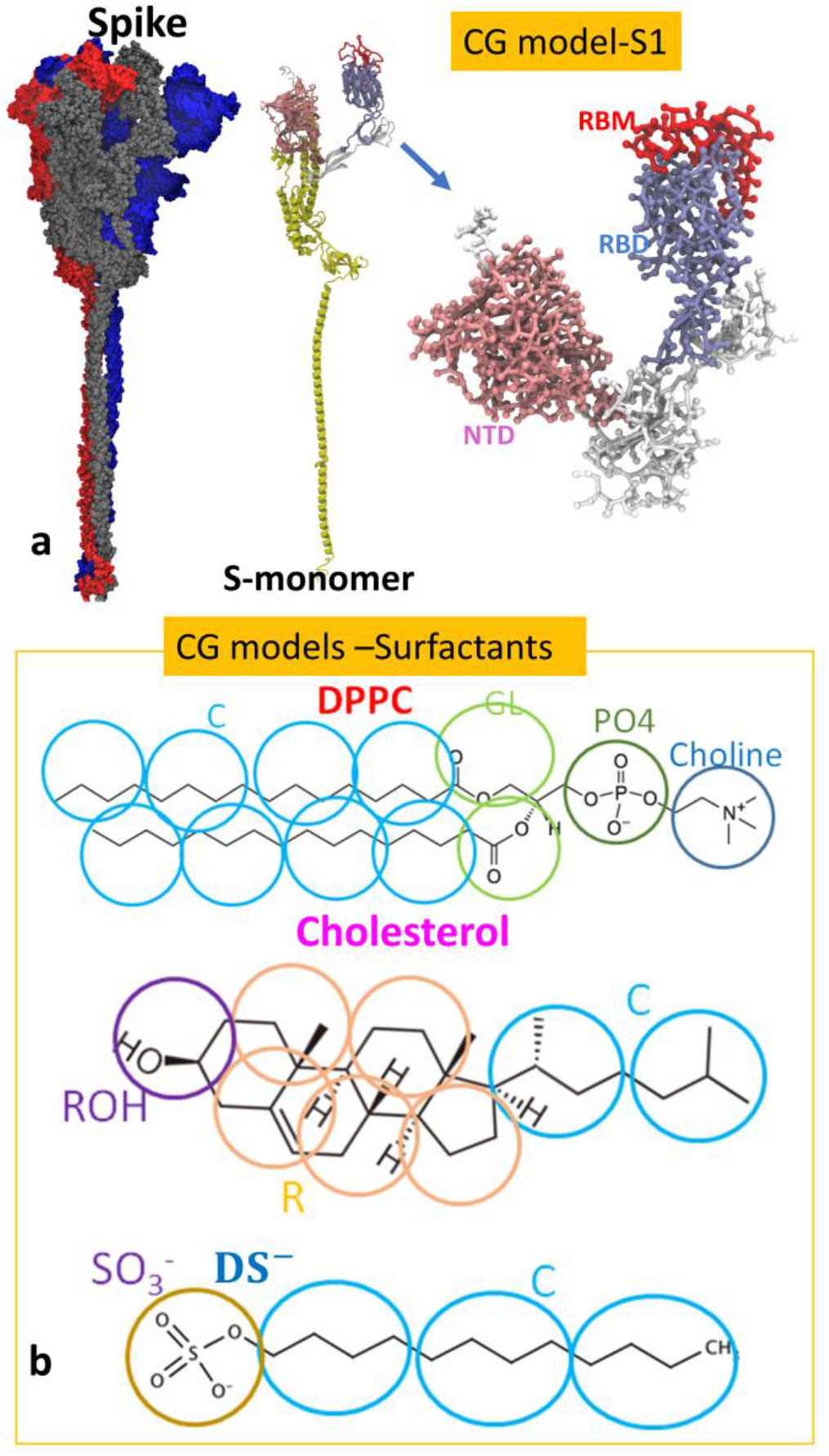
a) The atomistic models of Spike trimer and the monomeric S protein, and the MARTINI model of the S1 domain. NTD, RBD and RBM are colored respectively pink, iceblue and red. All other residues are shown in while. b) The MARTINI CG representation of the surfactants studied in the simulations.

**Table S1.**
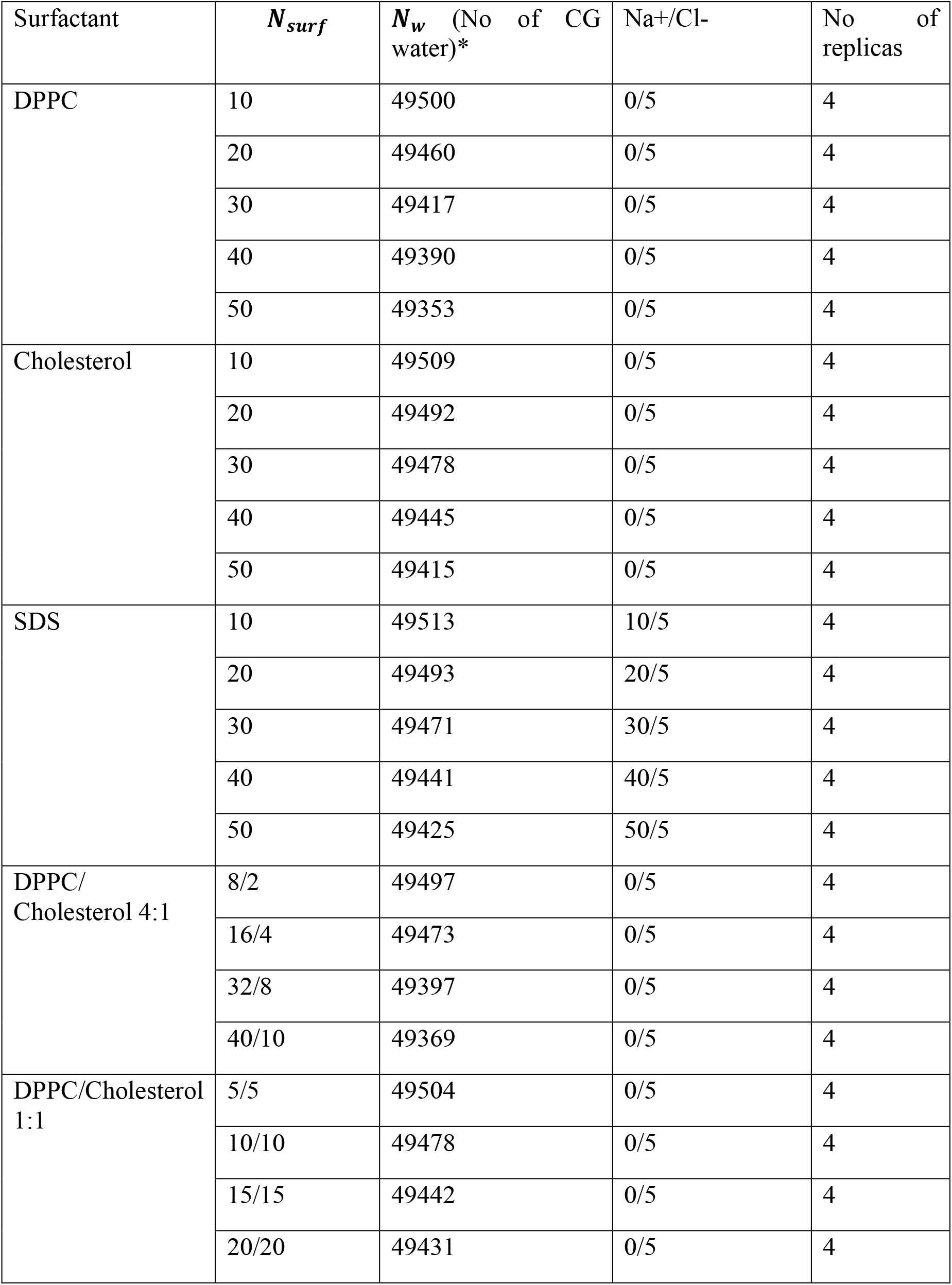
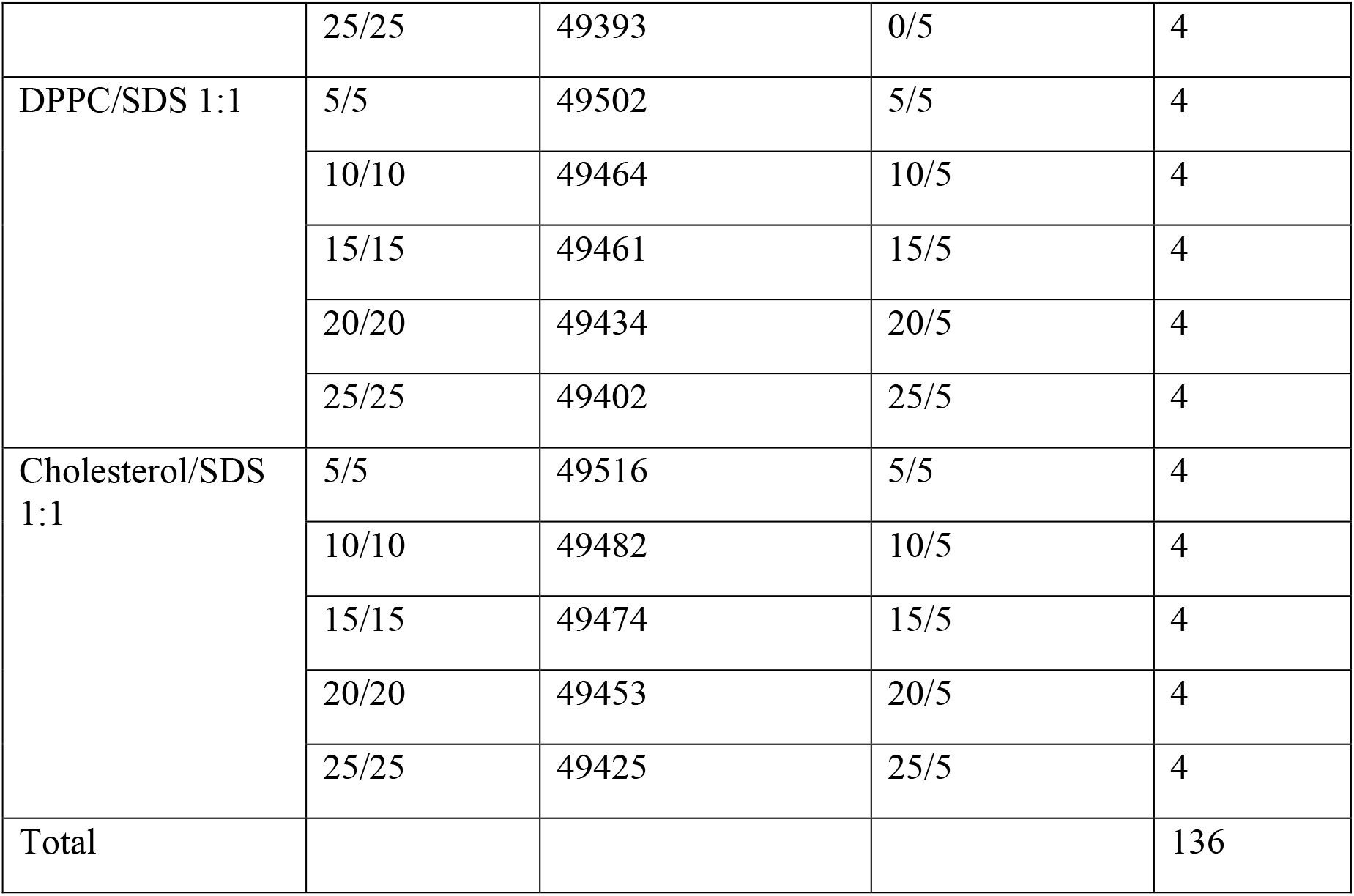
The details of the simulation systems. *The number of water CG beads given represents that in one of the replicas. N_w in the other three replicas may change by a few, typically less than 5.

#### SI. II Simulation protocol

MARTINI version 2^1–2^ is used for the simulations. The initial S1-surfactant systems are simulated a high temperature of 800 K with position restraints on the S1 fragment under NVT conditions, in order to randomize the surfactant distribution. The Coulomb interactions are incorporated with *reaction-field* scheme with a general non-bond interaction cutoff 1.1 nm and a dielectric constant *ε* = 15 that is appropriate for MARTINI standard water. Berendsen thermostat is used for temperature control and a time step of 20 fs is used. The final snapshot of this simulation is used as the initial configuration for the production runs. The initial configurations for the four replicas of each system are obtained with running the NVT randomization for different simulation times and with different random seeds for initial velocity generation; generally, NVT simulations are run in the range 0.2 -1.0 ns. Typical snapshots of initial configurations are provided in Figure S2. The production runs are performed for 1 *μs* for each system, at NPT conditions at physiological temperature 310 K and pressure 1 bar, controlled by Berendsen thermostat and barostat with a compressibility value 3 × 10^−4^ bar^−1^ and a coupling constant 12.0 ps.

**Figure S4.**
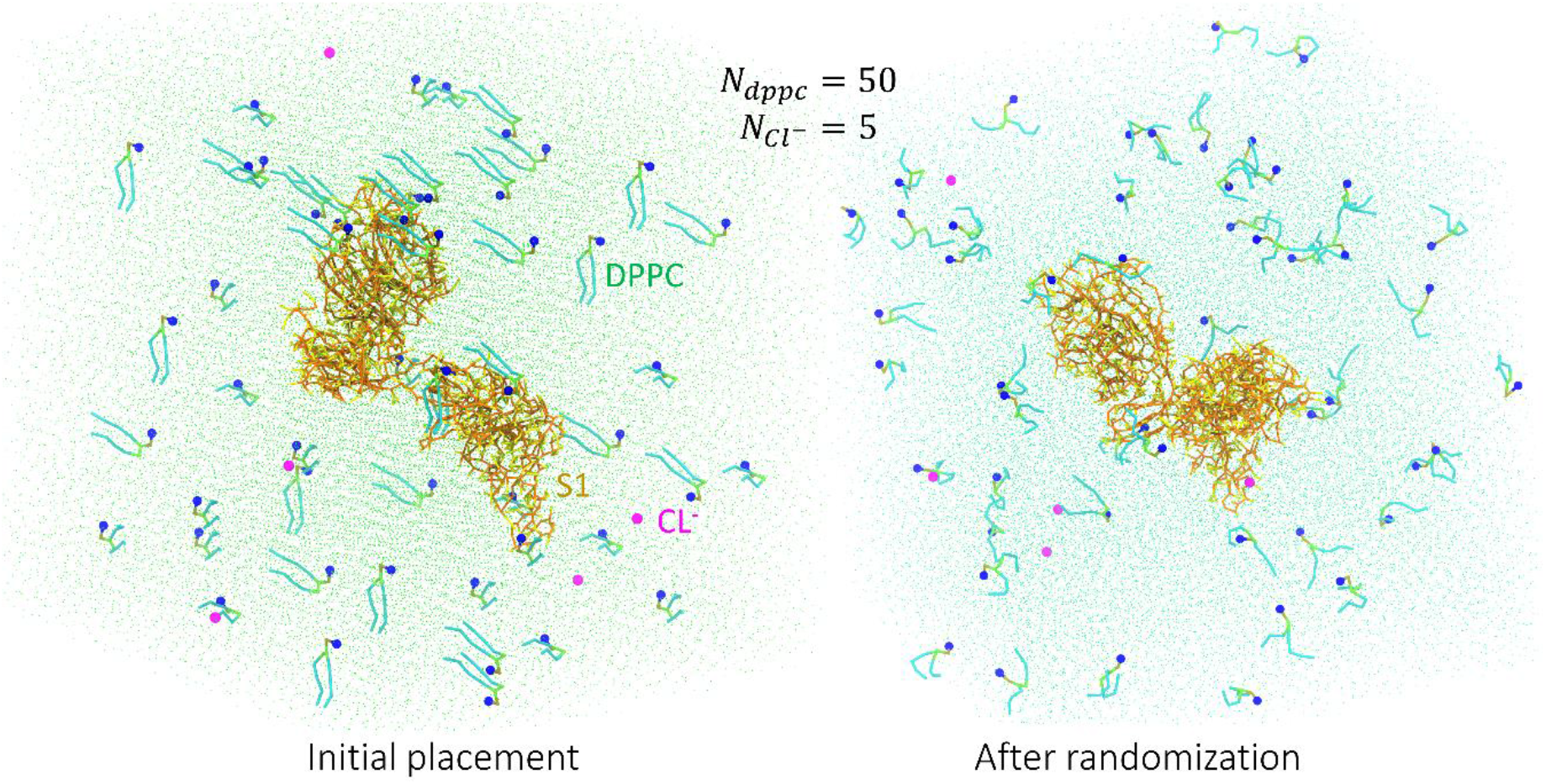
The snapshots of S1-DPPC system with *N_surf_* = 50 after the initial surfactant placement and solvation with water (left), and after randomization with NVT simulation at 800 K for 400 ps (right). Colors: S1 backbone is colored orange and the side chains in yellow, the choline group of DPPC in blue, the DPPC glycerol backbone in lime, DPPC tails in cyan, and the Cl^-^ ions in magenta. The water beads are shown as tiny green dots.

### SII. The surfactant adsorption probability of the S1 residues

The surfactant adsorption probability of the nth residue *P^ad^*(*n*) along the protein sequence is calculated as the fraction of time the residue is in contact with a lipid CG bead. The lipid contacts are counted when the distance between the protein and the lipid beads is less than or equal to 0.62 nm which is 1.3-1.4 of the LJ diameter of the MARITNI beads (0.47 nm). *P^ad^*(*n*) is calculated over 101 frames in the last 200 ns of each simulations. This is averaged over the 4 replicas at each surfactant concentrations, and further averaged over the 5 surfactant concentrations to finally obtain the values plotted in Figure 1b, Figure S3 and Figure S4. In Figures S3 & S4, the average probability in regions of relatively high (green) and low (red) adhesion are given. The highly adhesive regions are identified with relatively high probability of adhesion. The regions of residues,1-14, 100-150, 170-250, 320-506 and 530-600, are found to have high affinity to surfactants, in contrast to blocks 14-100, and 250-320, where surfactant adsorption is less probable. Note that similar patterns are observed in single component (Figure S3) and binary mixture (Figure S4) systems.

The average probability of an S1 residue to have adhered with a lipid bead in each simulation is calculated as,

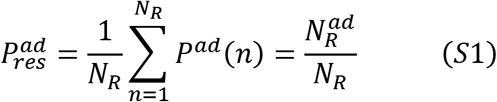

where *N_R_* = 685, the total number of residues in the S1 domain, and 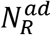 is the average number of residues adsorbed with surfactants.

**Figure S5.**
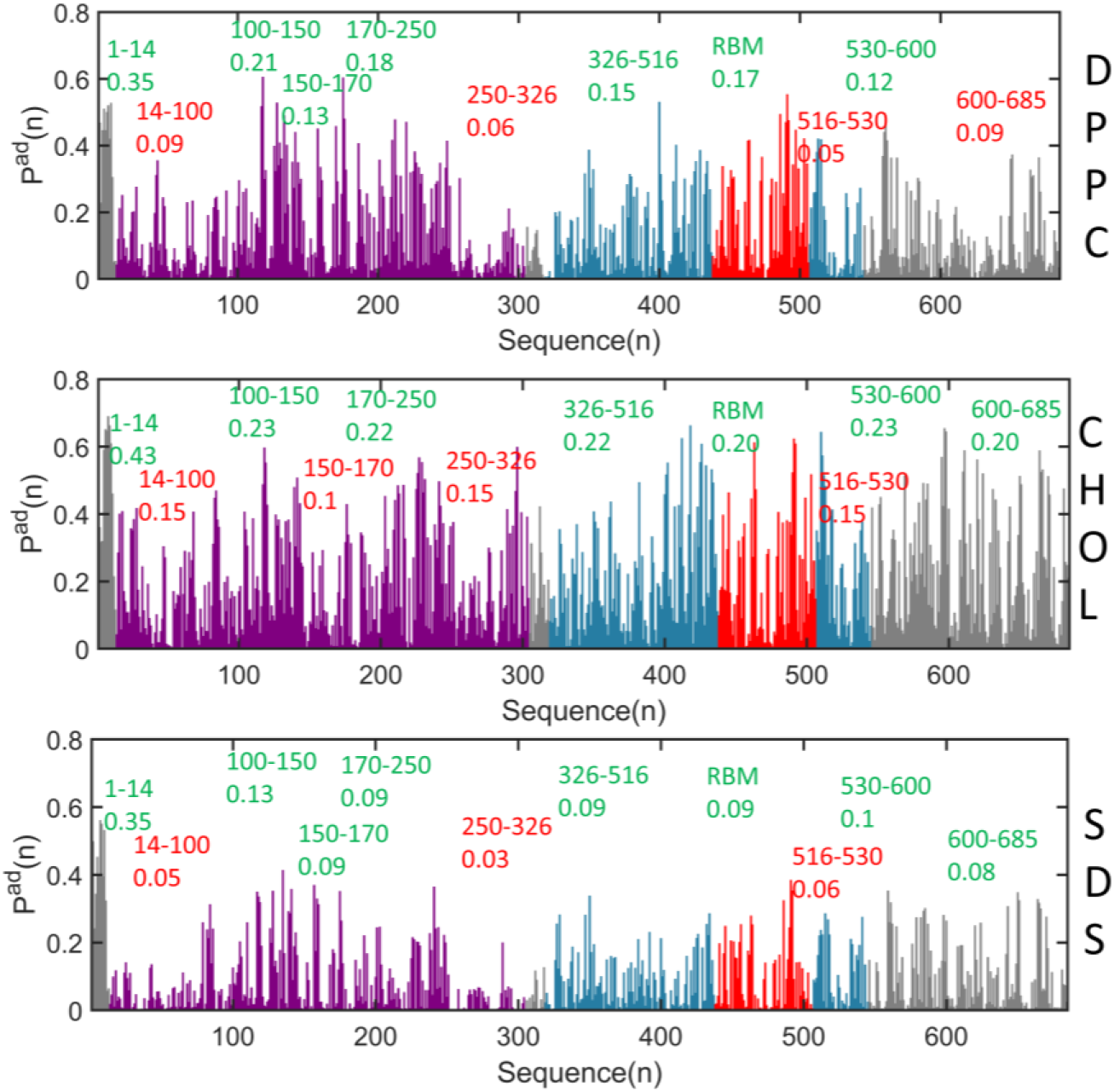
The surfactant adsorption probability of residues in the single componet surfactant systems. The residue regions with relatively high adsorption probability are indicated in green, while lower surfactant adsorption regions indicated in red. The average adsorption probabilities in those regions are also given. Other colors: purple-NTD, red-RBM, iceblue – RBD other than RBM and the remaining S1 regions – gray.

### SIII. Fraction of adsorbed surfactants and residue types

The fraction of surfactants obtained from the average number of surfactants in contact with the protein, 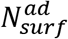 calculated in the simulation window 800-1000 ns over 101 frames as

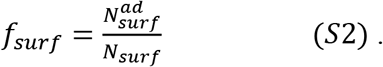

Note that, a surfactant may adhere to the protein, by its one or several CG beads. Thus a more quantitative picture will be obtained when the fraction of adhered surfactant CG beads are calculated, which is given by,

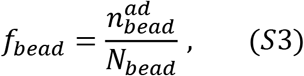

where 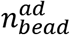 is the average surfactant CG beads in contact with the protein. The fraction of a particular bead type, such as choline or ROH etc is calculated in the same way. *f_bead_* in different cases are provided in Figure 3 of the paper.

**Figure S6.**
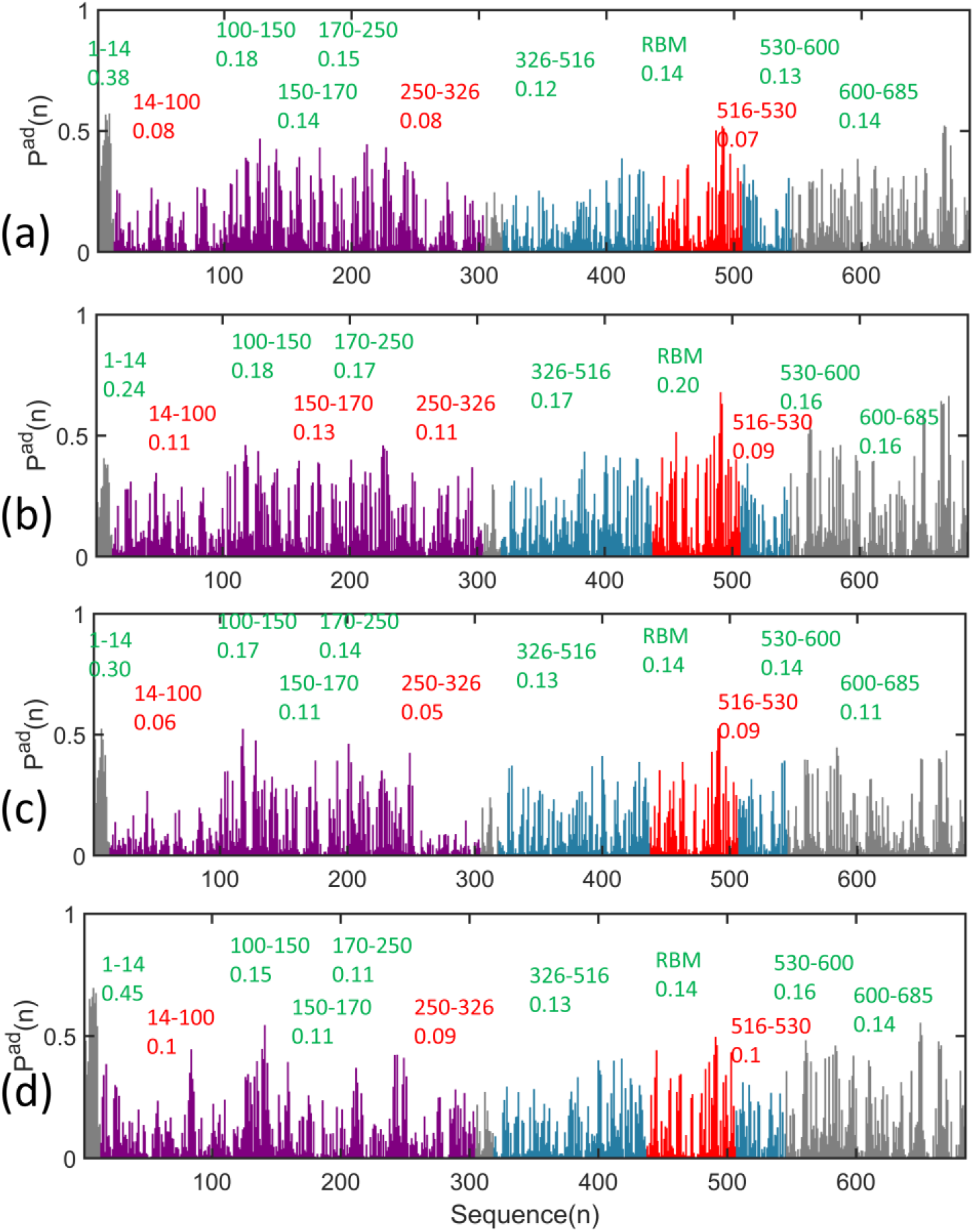
The surfactant adsorption probabilities in binary mixtures (a) DPPC:CHOL 4:1, (b) DPPC:CHOL 1:1, (c) DPPC:SDS 1:1 and (d) CHOL:SDS 1:1. The residue regions with relatively high adsorption probability are indicated in green, while lower surfactant adsorption regions indicated in red. The average adsorption probabilities in those regions are also given. Other colors: purple-NTD, red-RBM, iceblue – RBD other than RBM and the remaining S1 regions – gray.

The fraction of the surfactant adsorbed residue type,

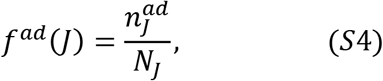

**Figure S5.**
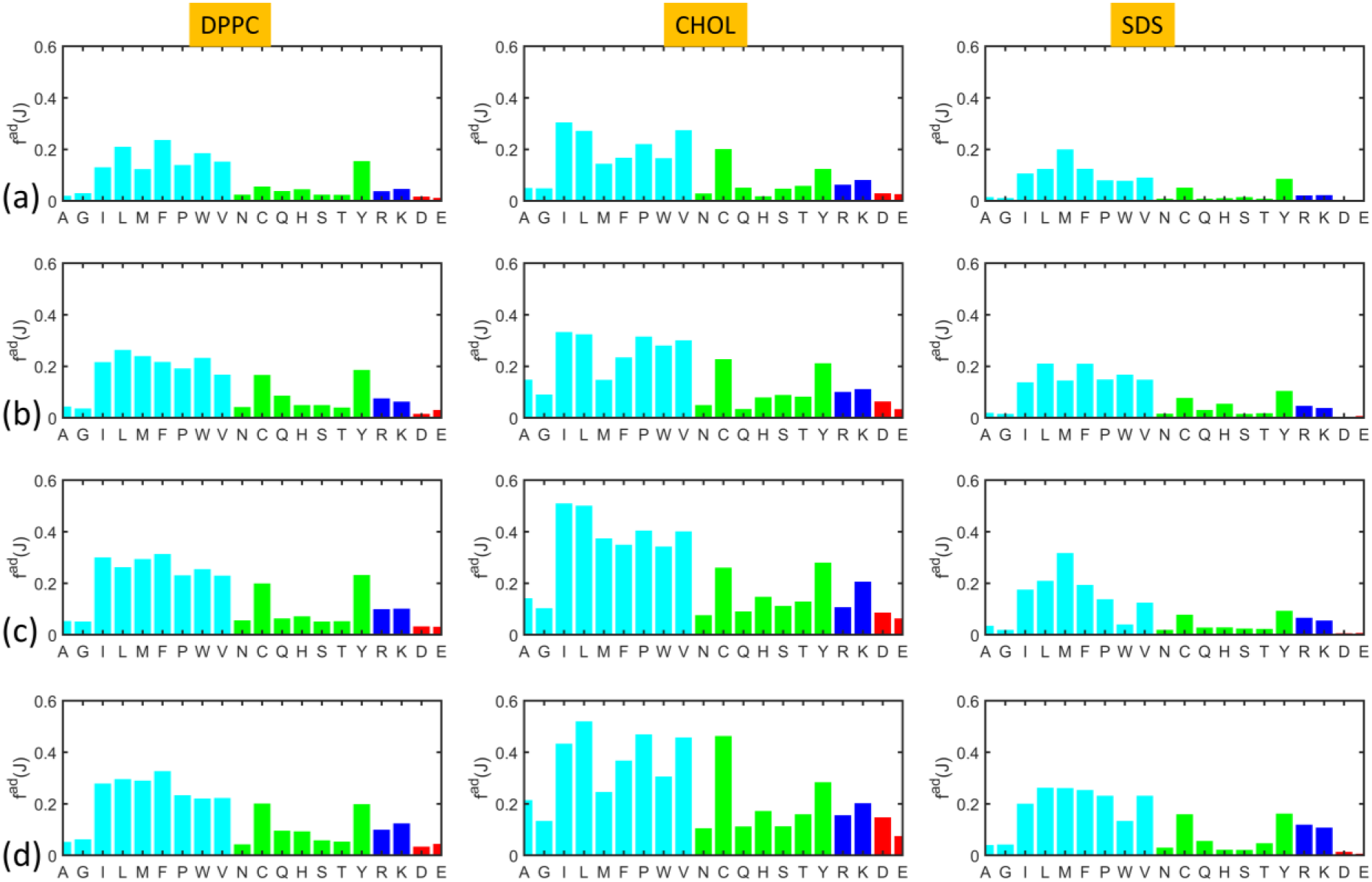
The fraction of surfactant adsorbed residue types in single-type surfactant systems at (a) *N_surf_* = 10, (b) *N_surf_* = 20, (c) *N_surf_* = 30 and *N_surf_* = 50. The hydrophobic, polar, cationic and anionic residues are respectively colored in cyan, green, blue and red.

where *J* = A, G, ,E, the amino acid types, 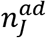 is the number of lipid-adsorbed amino acid residue of type *J* and *N_j_* is the total number of *J*-type amino acids in the S1 chain. *f^ad^*(*J*) for a given *N_surf_* is averaged over the four replicas. *f^ad^*(*J*) is plotted in the case of single type surfactant systems at various concentrations in Figure S5 and Figure 4b of the paper, which shows similar patterns for all surfactants at all concentrations. The hydrophobic residues ILE, LEU, PHE, PRO and VAL, and the polar CYS and TYR have pronounced affinity for surfactants. The cationic residues ARG and LYS are preferred by all surfactants, zwitterionic, anionic and non-ionic.

### SIV. Ionic residue substitutions in Delta and Omicron

Table S2 lists the ionic mutations in Delta and Omicron variants.^5–6^ Each variant is associated with an increase in ARG and LYS in the S1 domain.

**Table S2.** Analysis of ionic mutations in Delta and Omicron

| Variant | Mutations (S1-green, S2-red) | Net change in Ionic residues |
| --- | --- | --- |
| **Delta** | T19R, T95I, G142D, E156-, F157-, R158G, L452R, T478K, D614G, P681R, D950N | S1:<br>ARG +2; LYS +1<br>ASP 0; GLU -1<br>S2: ASP -1 |
| **Omicron** | A67V, del69-70, T95I, del142-144, Y145D, del211, L212I, ins214EPE, G339D, S371L, S373P, S375F, K417N, N440K, G446S, S477N, T478K, E484A, Q493R, G496S, Q498R, N501Y, Y505H, T547K, D614G, H655Y, N679K, P681H, N764K, D796Y, N856K, Q954H, N969K, L981F | S1:<br>ARG +2; LYS +3<br>ASP +1; GLU +1<br>S2:<br>ARG 0; LYS +3<br>ASP -1 |

